# Comparative analysis of the genomic architecture of six Fusarium species

**DOI:** 10.1101/2024.06.04.597288

**Authors:** Domenico Rau, Maria Leonarda Murgia, Davide Fois, Chiara M. Posadinu, Andrea Porceddu

## Abstract

Comparative analyses of several plant pathogens have revealed that genome plasticity could be associated with different genomic architectures. In certain species, genomic compartments are characterised by highly conserved regions that contain mainly housekeeping genes and rearranged regions that are enriched for genes related to virulence and adaptation. The compositional and structural characteristics of genomic regions have been significantly associated with compartment membership in single species, but little information is available on the covariation of these features between species.

Here, the results of a comparative analysis of the genomic architectures of six agriculturally relevant *Fusarium* species, which differ for several biological and pathogenic characteristics, are presented. These include *F. culmorum*, *F. fujikoroi*, *F. graminearum, F. oxysporum*, *F. solani,* and *F. verticillioides*.

The genome sequences of these species were partitioned into adjacent windows, with the average level of gene collinearity with the other species used as an index of compartment membership. High collinearity is typical of conserved regions, while low collinearity is typical of rearranged regions. Several genic and genomic variables were found to be consistently associated with compartment definition among all the *Fusarium* species that were investigated.

The compartment that was characterised by lower collinearity (i.e., high genomic rearrangements) contained more relocated genes, species-specific genes and secreted protein-encoding genes than regions with low collinearity. Furthermore, several molecular evidence indicates that low-collinearity regions are more likely to be subjected to selective pressure than high-collinearity regions. Indeed, genes residing in the former regions exhibited higher rates of sequence evolution than in the latter, as indicated by the high non-synonymous-to-synonymous substitution rates.

However, they exhibited signatures of selection to minimise the costs of transcription, as indicated by their high coding density. Our data suggests that although variable genomic compartments evolved mostly after species radiation, they share similar genomic features across related species and perhaps evolve with similar mechanisms.

## Introduction

The growing availability of complete genomic sequences of pathogens has facilitated our understanding of key aspects of genomic architectures associated with adaptability to new environments and/or hosts (Ma *et al*., 2013; Raffaele and Kamoun, 2012). Based on the distribution of DNA compositional properties and structural and functional features of genes, several types of genomic architectures of plant pathogens have been proposed (Frantzeskakis *et al*., 2019; Torres *et al*., 2020; Raffaele and Kamoun, 2012). One of the most prevalent and widespread genomic architectures is the so-called two-speed model, which posits that a mosaic genomic architecture is essential for the evolution of loci involved in virulence (Dong *et al*. 2015). The two-speed model is associated with the presence of two distinct genome compartments. The first comprises gene-dense and repeat-poor regions, which are home to essential and widely conserved housekeeping genes. The second is characterised by gene-sparse and repeat-rich regions, which contain fast-evolving virulence-associated genes (Dong *et al*., 2015; Faino *et al*., 2016). In addition, fast-evolving compartments are characterised by genomic regions rich in AT and the presence of accessory chromosomes (Yang *et al*., 2020; Bertazzoni *et al*., 2018).

The existence of subtypes of two-speed genomic architectures can be identified by examining the mode of compartmentalisation achieved (Möller and Stukenbrock, 2017; Priest *et al*., 2020). For example, in the *Verticillium dahliae* lineage, specific regions derive from duplications and rearrangements of effector genes flanked by transposable elements (Faino et al. 2016). In other instances, the presence of transposable elements can induce mutagenic mechanisms that can extend their effect to neighbouring effector genes (Frantzeskakis *et al*., 2019; Depotter *et al*., 2019). Species lacking a clear genomic compartmentalisation are considered to have a ‘one compartment’ genomic architecture. In these species, candidate secreted effector protein (CSEP) coding genes are not predominantly found in gene sparse areas as in other fungal pathogens (Frantzeskakis *et al*., 2018); it is believed that adaptation and virulence are governed by copy number variation (CNV) and heterozygosity of effector loci (Frantzeskakis *et al*. 2018).

The species belonging to the Fusarium genus exhibit a considerable degree of phenotypic, ecological, and genomic diversity (Ma et al. 2013). *F. verticillioides* is a pathogen that causes ear rot mainly in maize and sorghum, but it can also infect a wide range of other plant species. *F. solani* may behave as saprobes inhabiting agricultural and non-cultivated environments such as forests, littoral and coastal zones, and deserts (Mandeel et al., 2005), or as pathogens causing diseases on more than 100 plant species (Coleman et al., 2009). The complex of *F. oxysporum* species can cause wilt disease in more than 100 agronomically important plants. However, some isolates exhibit a high degree of host specificity and are classified as different formae speciales (Ma et al. 2013). *F. graminearum, F.culmorum*, along with *F. cerealis*, are the most prevalent pathogens that cause head blight in wheat and other cereals (Ma *et al*. 2013). *F.fujikuroi* is the pathogen that causes bakanae disease in rice plants (Ma *et al*. 2013).

The *Fusarium* spp phytopathogen can adopt a wide range of infection strategies. The majority of these fungi can be loosely classified as hemibiothrophs, as their initial infections resemble those of pathogens that depend on living hosts. However, they may eventually be considered necrotrophs, as the interaction can evolve to kill or consume host cells (Ma *et al*. 2013). Additionally, considerable variability has been observed in the reproduction mode. Some *Fusaria* produce meiotic (sexual) spores and up to three types of asexual spores. However, it should be noted that not all spore types are known to be produced by all species. In addition, fewer than 20% of *Fusaria* exhibit a sexual cycle. Some species are considered homothallic and/or homokaryotic, while others are heterotallic and/or heterokaryotic (Ma *et al*. 2013).

A common feature of *Fusarium* spp is the compartmentalisation of genomes into regions containing genes for essential functions (core genomes) and others containing genes for host specificity and pathogen virulence (adaptive/accessory genome) (Ma *et al*. 2013). However, the manner in which this compartmentalisation is achieved can vary between *Fusarium* species. For example, the four chromosomes of *F. graminearum* can be divided into two distinct regions: highly variable regions containing the majority of adaptive genes and less variable regions enriched for essential genes (Cuomo *et al*., 2007). Zhao et al. (2014) demonstrated that genes mapping in highly variable genomic regions of *F. graminearum* exhibit lower expression levels, higher sequence variability, and relaxed repeat-induced polymorphisms (RIP) compared to genes belonging to other genomic compartments (Zhao *et al*. 2014). These regions may function as genomic cradles, as genes that map therein after duplication undergo sub-functionalisation or even neo-functionalisation processes. Furthermore, these regions are also preferential sites for the integration of relocated or transposed genes. According to the authors, preferential relocation within a limited number of genomic cradles could lead to the formation of a cluster of functionally related genes (Zhao *et al*. 2014). Other species, such as *F. oxysporum*, *F. solani,* and *F. fujikoroi,* exhibit supernumerary chromosomes whose number and gene content are highly variable within the species (Bertazzoni *et al*. 2018). These species also contain variable regions within core chromosomes that show features typically associated with accessory chromosomes and have been designated Accessory Regions (ARs). ARs have been shown to be dispensable (Bertazzoni *et al.,* 2018).

The compositional and structural characteristics of genomic regions have been significantly associated with compartment membership in a single *Fusarium* species. However, little information is available on the covariation of these features between species. The identification of trends shared between species or unique to a single species is expected to provide information on mechanisms of adaptive evolution, also with respect to host preference. For example, when comparing *F. oxysporum* strains isolated from human and plant-infected tissues, although the overall structure of the chromosome was similar, some features were found to be markedly different (Zhang et al. 2020). The accessory chromosomes in human isolates were found to be enriched in genes encoding proteins for the transport of metal ions and inorganic cations, while those of phytopathogens were found to be enriched in plant cell wall-degrading enzymes (Zhang et al. 2020). Furthermore, the transposable elements present in the two strains exhibited substantial differences, although their role in promoting sequence evolution and rearrangements was likely comparable (Zhang et al. 2020).

In the present study, we examined genomic regions of six Fusarium species that differ for several biological and pathogenic characteristics. These include namely *F. graminearum*, *F. culmorum*, *F. fujikoroi*, *F. oxysporum*, *F. verticillioides,* and *F. solani*. The genome sequences of these species were divided into regions with high (conserved) or low (rearranged) gene collinearity. The compositional variables of these regions and the structural characteristics of the genes residing within them were then associated with the level of collinearity and compared across species.

## Materials and Methods

### Sequence and annotation datasets

Genomic sequences and annotations were obtained from the Ensemble database (https://fungi.ensembl.org/). For *F. graminearum,* the genomic sequence of the PH1 strain was obtained from the Rothamsted Centre for Genome Analysis (accession PRJEB5475), deposited as the assembly GCA_900044135.1 in the International Nucleotide Sequence Database Consortium. For *F. culmorum*, we used the assembly of chromosomes 1-4 deposited as the assembly GCA_900074845.1 (assembly name FcUK99v1.2). Contigs 5 and 6 were not considered since they are assembled without physical map evidence.

For *F. fujikoroi,* the genome build accession GCA_000315255.1 from the Joint Genome Institute EF 1 (genome version EF1) was considered. The genome sequence, assembly, and annotation of protein-coding genes of the *F. oxysporum* genome were generated by the Broad Institute as part of their work on the Fusarium Comparative Sequencing Project (genome-build GCA_000222805.1, genome version PO2). The assembly and annotation of *F. solani* used in this study were obtained for JGI (V2.0 INSDC assembly GCA_000151355.1). The assembly and annotation of *F. verticillioides* were provided by the Broad Institute (genome build GCA_000149555.1).

Genomic sequences were subjected to a series of analyses that included extraction of exons, introns, and intergenic sequences by using gff2sequence software (Camiolo and Porceddu 2013).

### Synteny among chromosomes and duplicate classification

Sequence duplicates were classified using MCScanX (Wang *et al*. 2012) using default settings. Briefly, all query proteins of each species were compared with proteins of the other investigated species and with themselves using BlastP and the best five non-self-matches with an e-value below 1e^-10^ were reported. The resulting hits were studied according to the position of the genes on the chromosomes and scaffolds. The highest scoring path was identified by dynamic programming as implemented in McScanX with standard settings (-k 50; -g -−; -s 5; -m 25). The Index of Local Collinearity (ILC) was defined as the proportion of loci mapping within a 100 kb window that are collinear with another species. Genomic regions were assigned to genome compartments based on the latent hidden Markov model calculated with depmixS4 (Visser and Speekenbrink 2010). A two-state model was considered for *F. graminearum, F. culmorum,* and *F. verticillioides,* while a three-state model was considered for *F. fujikoroi, F. oxysporum, and F. solani* to account for the presence of accessory chromosomes.

### Identification and masking of repetitive sequence

Repetitive sequences were identified using the RepeatModeler2 (Flynn *et al*. 2020) pipeline with the default settings. The identified libraries were subsequently used by RepeatMasker (Smit *et al*. 2013) to mask repetitive elements within genomes.

### Pseudogene identification

Pseudogenes were identified using a homologous search approach as implemented in Exonerate software (Slater and Birney, 2005). In summary, full-length protein sequences were used as queries against the subject genome, which had previously been masked to remove repetitive and genic regions. The output file was formatted in accordance with the specifications indicated in the command line %qi\t%ti\t%pi\t$em\t%ps\t%qab\t%qae\t%tab\t%tae\t%r\t%s\n.

Alignments with greater than 40% sequence identity were retained for subsequent analyses (Mascagni *et al*. 2021). If a genomic region exhibited sequence similarity to multiple query proteins, the optimal pseudogene query pair was identified based on the Exonerate score and alignment identity.

### Bioinformatic identification of secretomes

The prediction of secretomes in the five *Fusarium* spp. was based on the protocol described by Brown et al. (Brown *et al*. 2012), with minor modifications. All protein sequences were initially analysed with SignalP (Almagro Armenteros *et al*. 2019) and those predicted to have a signal peptide were filtered and analysed with TargetP (Emanuelsson *et al*. 2007). The sequences identified as having a signal peptide by TargetP were scanned for the transmembrane domain using TMHMM (Krogh *et al.,* 2001) and all sequences with 0 or 1TM, if located in the signal peptide, were retained. The software PROTCOMP (Klee and Ellis, 2005) was also employed to predict the localisation of the sequences already filtered, as previously described. All proteins predicted as extracellular or unknown were subjected to further analysis using the WolfPsortII package (Horton et al., 2007) in order to obtain the final data sets of secreted sequences.

### Analyses of duplicate depth, relocated, and species-specific genes

The presence of gene homologues within each species was investigated by ‘all *versus* all’ BLASTP searches (Altschul *et al*. 1990). Pairwise alignments that showed identity greater than 50% and coverage greater than half the length of the shortest protein in the alignment were filtered. The BLASTP output was parsed with the OrthoMCLBlastParser.pl script v2.0 (Chen *et al*., 2006).

The duplication depth (DD) of a gene was calculated by considering the number of parsed alignments. For example, the DD of a given gene was equal to 1 if it showed only self-hit or greater than 1 if it showed additional hits with identity >=50% and alignment coverage >=50%. The aDD was calculated by averaging the DD of all genes within a given 100 kb region.

To identify species-specific genes, all versus all BLASTP searches were performed and queries that showed no hits to sequences of other species were selected.

Relocated genes were identified as loci that lacked both homologous genes and collinear orthologous genes in other species (Hart *et al*. 2018a). In this analysis, the threshold for identifying a sequence as homologous to a given locus was established at 50% of the amino acid sequence identity, calculated on alignments of at least half the length of the aligned proteins.

### Evolutionary rate calculation and codon usage

Orthologous groups were identified using Orthofinder software (Emms and Kelly 2019) with default settings. The codon alignments were obtained by passing the protein alignment obtained with Clustalo (Sievers *et al*. 2011) and the nucleotide coding sequences to the PRANK software (Löytynoja and Goldman 2010) with the default settings.

The synonymous substitution rates (Ks) and non-synonymous substitution rates and the omega ratios (Ka/Ks) were determined using the yn00 software in the PAML package (Yang 2007). For each gene, the averages for Ka, Ks and the omega ratio calculated from pairwise alignments with orthologs from other species were considered. The codon bias was estimated using the effective number of codons as implemented in the software ENC with default settings (Novembre 2002).

### Identification of repeat-induced point mutations

Repeat-induced polymorphism (RIP) is a mechanism that has been postulated to be evolved to inactivate repeat elements in the fungal genome by introducing C-to-T transitions in duplicated sequences (Galagan and Selker, 2004). *In silico* investigations of RIP are based on dinucleotide frequency in genomic windows. In this study, three compositional indices were employed. The product index is defined as the ratio between TA and AT in windows of 1000 bp. The substrate index is calculated as the ratio between (CpA + TpG) and (ApC + GpT). The composite index is the difference between the product index and the substrate index. These RIP indexes were calculated using RIPPER web tools with useful parameters (Van Wyk *et al.,* 2019). A region is considered to be affected by RIP when its RIP composite index exceeds 1.1, the RIP substrate index is below 0.9, and the RIP composite index is above 0 (Van Wyk *et al.,* 2019).

### Analysis of compositional (regional) variables

This involved determining the mononucleotide and dinucleotide content of each genomic region was determined using Perl scripts developed in-house. Dinucleotide bias was calculated as the ratio between the relative frequency of a given dinucleotide and the product of the composing mononucleotide (Karlin, and Ladunga, 1994; Karlin, 1998). The relative abundance difference between pairs of regions was calculated as the mean of the absolute difference between the dinucleotide biases of the analysed pair (Karlin, and Ladunga, 1994; Karlin 1998).

### Analysis of regional structural variables

The gene content of each 100kb region was determined using the intersectBed application of Bedtools (Quinlan and Hall, 2010). For each 100 kb region, the gene content was calculated as the number of genes mapped within the region. In the event that a gene overlapped the boundary between two regions, the degree of membership in either region was proportional to the length of the portion of the overlapping gene. The content of secreted and species-specific genes was determined by calculating the ratio between the number of these genes (identified as explained above) and the total number of genes in the region. The content of repetitive elements and pseudogenes was calculated as the proportion of each 100-kb region classified as repetitive or pseudogene, respectively.

### Analysis of gene composition and structure

This involved determining the structural parameters of the genes, which were based on the gff3 files of each species. The dinucleotide compositions of DNA sequences were analysed in accordance with the methodology proposed by Karlin and Burge (1995).

### Circos graphs

A comprehensive representation of the genomic landscape was produced for each species using Circos version 0.69-9. (Krzywinski *et al.,* 2009).

### Statistical analysis

The association between variables and compartment membership was analysed using the Wilcoxon non-parametric test as implemented in R. The significance of Wilcoxon tests was also tested against a null distribution obtained by randomly permuting either genes or compartment membership among regions for genic and regional variables, respectively. Principal component analysis (PCA) was carried out with the R fviz package and visualised with the R factor extra (R Core Team 2013).

## Results

The genomic compartments of six species of *Fusarium* (namely *F. culmorum, F. graminearum, F. fujikoroi*, *F. oxysporum*, *F. verticillioides* and *F. solani*) were identified by collinearity analysis (Zhao *et al*., 2014; Hane *et al*., 2011).

For each 100 kb adjacent genomic region, the Index of Local Collinearity (ILC) was defined as the proportion of the total number of loci mapping within the region and classified as collinear to loci of another species. Genomic regions exhibiting high ILCs were assigned to the genomic compartment designated as ‘A’. Those exhibiting intermediate ILCs were assigned to the compartment designated as ‘B’. The compartment designated as ‘C’ included all the regions of the accessory chromosomes in addition to a subset of regions of the *core* chromosomes exhibiting an ILC of 0 or nearly 0 (Figure 1 and Supplemental Figures S1-S4). The membership of each 100 kb window to the genomic compartment is presented in Supplemental Table S1.

**Figure 1.**
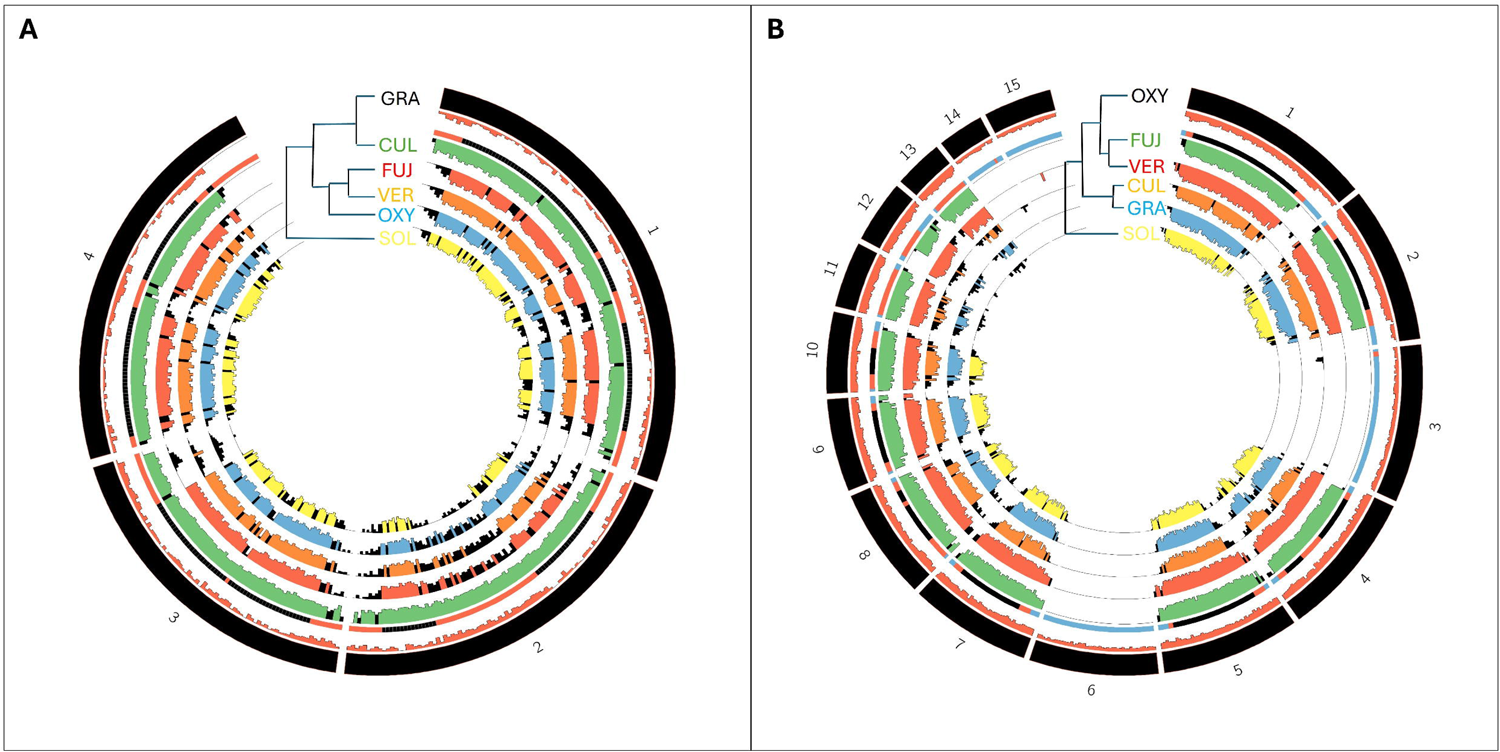
Circos plots illustrating trends for the index of local collinearity (ILC) and the definition of genomic compartments. For each circos plot (A and B), proceeding inward, the first whorl (in black) is an ideogram of chromosomes of the investigated species, *F. graminearum* and *oxysporum*, respectively; in the second whorl, histograms (in dark Orange) represent the gene counts within each window of 100kb. The tiles of the third whorl represent genomic compartments: black = compartment A, red = compartment B, blue = compartment C. The other whorls represent ILC trends with the other species. The phylogenetic tree of the investigated species indicates also the species used to calculate ILC. Abbreviations of species names are: CUL for *Fusarium culmorum*; FUJ for *F. fujikoroi*; GRA for *F. graminearum*; OXY for *F. oxysporum*; SOL for *F. solani*; and VER for *F. verticilliodies*.

Three main trends were evident. First, pairs of species with low phylogenetic distances showed higher average ILC values. For example, when comparing the recently diverged *F. graminearum* and *F. culmorum*, the ILC values for compartments A and B were 0.81 and 0.70, respectively. However, when comparing *F. culmorum* and *F. solani*, which have a more ancient divergence, the ILC values for compartments A and B dropped to 0.656 and 0.132, respectively (Figure 1A). Second, for all species under analysis, the genomic regions with low (≤ 0.5) or high (≥ 0.5) ILC values were observed to cluster in the core chromosomes. Third, an ILC value of 0 was observed indicating that there was no collinearity for the accessory chromosomes of *F. oxysporum* (Figure 1B), *F. fujikoroi* (Figure S2) and *F. solani* (Figure S3), as well as for a few regions of the core chromosomes, which have already been defined as accessory regions (Bertazzoni *et al*., 2018).

### Structural and compositional variables are associated with compartment membership

Each 100 kb genomic region was analyses for several compositional and structural characteristics, which we subsequently designated as ‘regional’ (genomic) variables.

The number of genes within each 100 kb region (regional gene density: rgd) was found to be a clear distinguishing factor between genomic compartments. On average, rgd was significantly higher (P < 0.01; Wilcoxon test) in compartment B than in compartment A in all species analysed (Figure 2A). Even the lowest difference observed between compartments A and B of *F. culmorum* (32.18 versus 33.77) was confirmed by a permutation test conducted by shuffling the memberships to compartments of the regions (Table S2A). The genomic regions within compartment C showed the lowest average rgd in *F. oxysporum* (17.95) and *F. fujikoroi* (24.71). In *F. solani,* chromosomes 14, 15 and 17 showed a low rgd (22.30, 17.75 and 19.0, respectively), in accordance with expectations for accessory chromosomes (see supplementary Figure S5E). In contrast, chromosomes 7, 12, and 16, despite being also assigned to compartment C, showed an unusually high gene density (35.37, 36.25 and 25.0, respectively), similar to that observed in core chromosomes (see supplementary Figure S5E).

**Figure 2.**
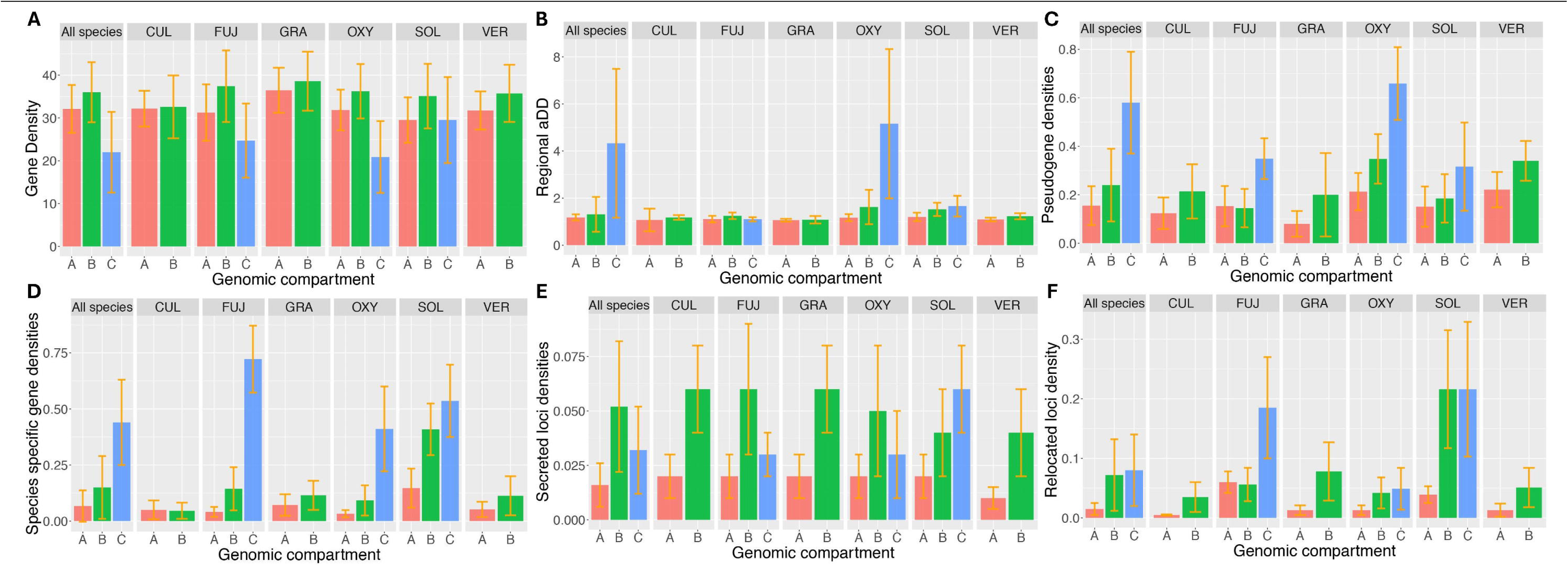
Association between genomic compartments and structural regional variables: gene density (A), regional aDD (B), pseudogene densities (C), species specific gene densities (D), secreted loci densities (E) and relocated loci density (F). Compartments A and B includes regions with high and intermediate ILC, respectively, while compartment C includes regions with ILC = 0 (see text). Abbreviations of species names are: CUL for *Fusarium culmorum*; FUJ for *F. fujikoroi*; GRA for *F. graminearum*; OXY for *F. oxysporum*; SOL for *F. solani*; and VER for *F. verticilliodies*. All species reports histograms of the complete dataset which included all analyzed species.

The repeat content (rc) (as bp of each region covered by repeats) did not show a consistent pattern across species. The mean rc was higher in the compartment with low ILC (i.e. compartment B and C) than with high ILC (i.e. compartment A) in the Fusarium species with accessory chromosomes. As an example, the mean rc of the *F. oxysporum* regions belonging to compartments C and A were 36945.00 and 1621.05 respectively, with a highly significant difference (P value <0.001). However, the difference in rc between compartments was less pronounced in the other species, with the exception of *F. culmorum* which showed a higher rc value in compartment B (3705.49) than in compartment A (1472.93). The full spectrum of rc differences between compartments is reported in Table S2B.

The duplication depth (DD) of each locus was calculated as the number of loci that show sequence similarity above a specified threshold. Sequence thresholds employed were at the protein level, as detailed in MM). In all species analysed, the majority of loci did not align with other loci and were classified as singletons (DD = 1). The species *F. graminearum* showed the highest percentage of singletons (90.4%). The lowest percentages were observed in *F. oxysporum* and *F. solani* with 73.8% and 79.9 %, respectively (Supplemental Table S3).

The average (regional) duplication depth (aDD) was found to discriminated between genomic compartments in all species investigated (see Figure 2B). Indeed, higher aDD values were observed in regions assigned to compartments B and C (1.31 and 4.33) than in compartment A (1.11). The only exception to the general trend was observed in *F. fujikoroi,* for which the aDD of compartment C showed lower values (1.1) than of compartment B (1.24) (Figure 2B; see also Supplementary Table S2C for results of the permutation test).

The distribution of duplicated genes and of the total number of functional genes could be influenced by intragenomic variations in sequence duplication and/or decay (Cole et al., 2001). For example, some chromosome regions may have experienced higher rates of gene inactivation subsequent to duplication than others, resulting in a relatively lower number of genes compared to the number of pseudogenes (Mascagni et al., 2021). The intergenic regions of the six Fusarium spp. were analysed for sequences showing homology to the predicted functional proteins, resulting in 33,934 regions (manuscript in preparation). For each genomic region, the number of pseudogenes was calculated as a proportion of the total number of loci (both functional and pseudo) mapping therein. In all investigated species, there was a moderate effect of compartment membership on pseudogene density. Regions with low ILC exhibited a significantly higher abundance of pseudogenes than other regions (Figure 2C and Table S2D).

#### Chromosome regions with low ILC exhibit higher density of species-specific genes

A comparative analysis of proteome sequence diversity revealed that a significant number of loci lacked homologs in other *Fusarium* species (Ma *et al*. 2013). We defined species-specific genes as those with no homologous counterparts in the other species analysed (see Table S3 for the lists of these loci). The species *F. solani* had the highest number of species-specific genes (3949 corresponding to 25.1% of all genes), followed by *F. oxysporum* (*2368*; 14.0%) and the remaining four species, namely F. *fujikoroi* (1221; 14.0%), *F. verticillioides*: (1,182, 8.3%), *F. graminearum* (1179; 8.3%), and *F. culmorum* (709, 5.6%). In all species investigated except *F. culmorum*, species-specific genes were found more frequently in regions of compartments B and C than in regions of compartment A (Figure 2D and Table S2E). A comparable picture was observed for the frequency of genes lacking orthologs in the other species, as defined by a reciprocal best blast score implemented in OrthoMCL (Chen et al., 2006) (see Figure S6).

#### Chromosome regions with low ILC have a higher density of secreted and relocated genes

In all species analysed, the proportion of genes encoding secreted proteins was higher within regions characterised by low ILC than in other regions (compartments B and C in Figure 2E and Table S2F). None of the 1,000 replicates generated by shuffling gene positions between genomic regions could reproduce the observed order of disproportion (i.e., P < 10^-3^). The complete list of loci identified as putatively secreted in the analysed *Fusarium* spp. is reported in Table S3.

A gene was defined as a relocated gene if it was a singleton (DD = 1) and its orthologs were not collinear (Hart et al., 2018b). These genes may have arisen from the duplication of an existing gene that was then inactivated, or from the transposition of a gene to a new genomic position (Hart *et al*., 2018). A significant association was identified between the content of the relocated genes within each 100 kb region (number of relocated genes / total number of genes mapping in the region) and the membership to the compartments (Figure 2F). Indeed, on average, a higher proportion of relocated genes was observed in the B and C compartments compared to the A compartment (Figure 2F and Table S2G). The lists of relocated genes in the Fusarium spp. are presented in Table S3.

#### The DNA composition of the low-ILC chromosome regions is A+T rich

A bias toward a high A+T content was observed in regions with low ILC in all species studied (A = 0.5023, s.d.= 0.024; B = 0.49, s.d.= 0.13 and C = 0.48 s.d. = 0.09; Figure S7). In addition, the frequency of several dinucleotides was found to vary significantly between genomic compartments (Figure S8). We then sought to determine whether the two compartments had different compositional biases. To this end, we considered the delta distance between regions, which is the difference between the observed dinucleotide frequency and that expected based on mononucleotide composition (Karlin and Burge, 1995). For all species analysed except *F. solani,* the delta values were found to be significantly higher for comparisons involving regions belonging to different genomic compartments than for pairs of regions that were randomly sampled regardless of compartment membership (P < 0.01 and Figure S9).

### Structural and compositional properties of genes and genomic compartment’ membership

Genomic regions were also characterised based on the compositional and structural features of the genes mapping within them (Figure 3). The regional value of a given genic variable was determined by averaging the values of that variable for all genes mapped within the region under consideration. Several regional variables were found to be associated with compartment definition (Figure 3 and Table S2H-O). The regions assigned to compartment A had, on average, longer coding sequences than those assigned to compartments B and C (Figure 3A and Table S2H). These associations were consistently observed in all species examined, with a general trend of A > B > C (Figure 3 and Table S2H).

**Figure 3.**
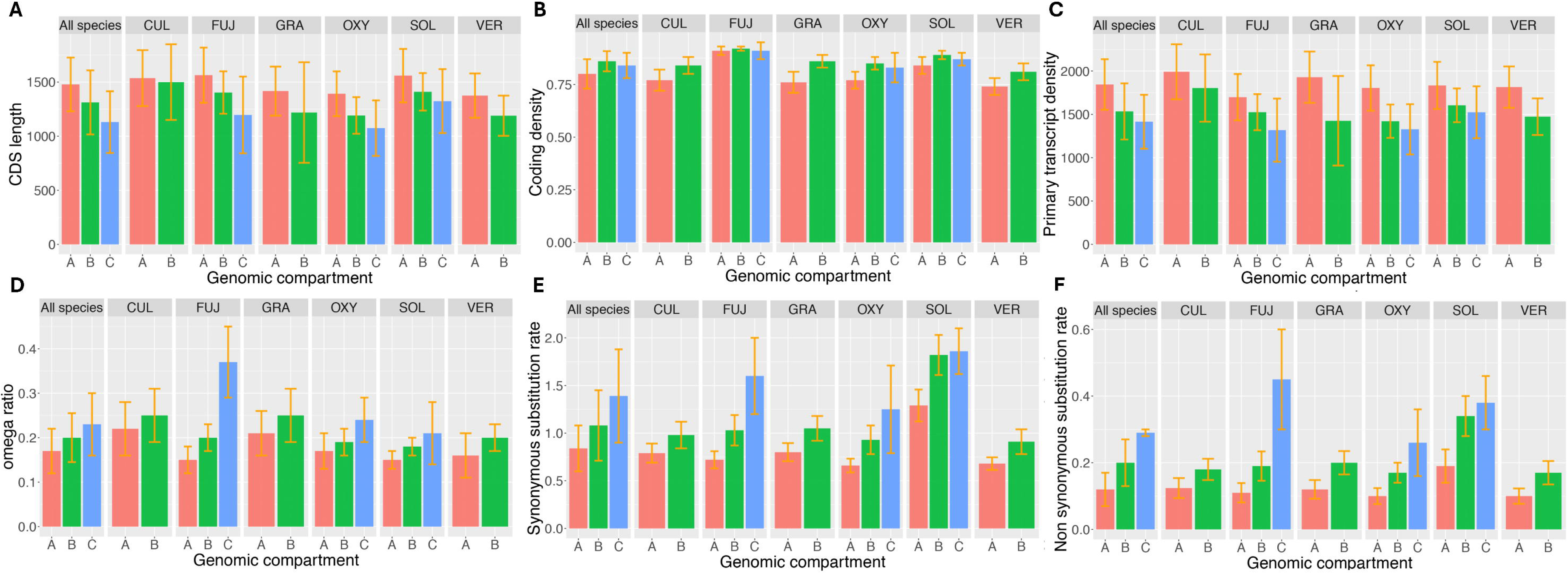
Association between genomic compartments and regional genic variables: CDS length (A), coding density (B), primary transcript density (C), omega ratio (D), synonymous substitution rate (E), non synonymous substitution rate (F). Compartments A and B includes regions with high and intermediate ILC, respectively, while compartment C includes regions with ILC = 0 (see text). Abbreviations of species names are: CUL for *Fusarium culmorum*; FUJ for *F. fujikoroi*; GRA for *F. graminearum*; OXY for *F. oxysporum*; SOL for *F. solani*; and VER for *F. verticilliodies*. All species reports histograms of the complete dataset which included all analyzed species.

The coding density (i.e., cds length / total primary transcript length) was higher in regions of compartments B and C than in regions of compartment A. The statistical robustness of all these observations was confirmed by the permutation test (P<0.01). For a comprehensive picture of the genic variables analysed, see Tables S2H-J. It should be noted that primary transcripts (i.e. exons plus introns) were significantly longer in compartment A compared to other regions (Figure 3C and Table S2J).

The ratios between non-synonymous / synonymous substitution rates (omega ratios) of genes were obtained by averaging the substitution rates calculated from pairwise alignments of orthologous coding sequences. The omega ratios were significantly (P < 0.01) higher for the regions of compartments B and C than for compartment A, indicating an enrichment of nonsynonymous substitutions in the first two compartments (Figure 3F). This pattern was also observed for both the synonymous (Ka) and nonsynonymous (Ks) substitution rates separately (Figure 3D and 3E, respectively), i.e. regions of compartments B and C are tout-court more prone to mutation than compartment A, with a higher bias towards nonsynonymous mutations.

The effective number of codons (ENC) is a synthetic measure of codon bias that can vary between 21 (maximum bias) and 59 (no bias) (Novembre, 2002). The genomic regions belonging to compartments B (ENC= 54.24) and C (ENC=54.24) showed significantly lower codon bias (P<0.01) than those of compartment A (ENC=53.4).

Repeat-induced point mutation (RIP) is a mutagenic mechanism that acts against repetitive elements (Galagan and Selker, 2004). *Fusarium* genomic sequences were analysed with the RIPPER software (Van Wyk *et al*., 2019) in order to calculate the compositional index associated with the RIP activity.

Approximately 5% of the genome of *F. solani*, *F. culmorum,* and *F. fujikoroi* showed RIP indexes (see Materials and Methods) indicative of the occurrence of RIP (Table S4). In contrast, the proportion of the genome affected by RIP was less than 1% in *F. graminearum*, *F. oxysporum* and *F. verticilliodes* (Table S4). The distribution of RIPPED regions among the genomic compartments was then analysed. A significant association (P < 0.01) was found between the genomic compartments and the frequency of 100 Kbp genomic windows that showed at least one evidence of repeat-induced mutations in *F. graminearum*, *F. solani*, *F. culmorum* and *F. oxysporum* (Table S4). In particular, the genomic windows of compartments B and C exhibited regions with a higher RIP frequency than would be expected by chance. Conversely, no significant associations were observed for *F. fujikoroi* and *F. verticilliodes*.

### Contribution of regional and genic variables to compartment differentiation

Principal component analysis (PCA) was employed to investigate the pattern of differentiation among the genomic regions under consideration. First, all species and all variables (regional and genic) were included in the analysis (Figure 4). The distinction between species was more evident than between compartments (see Figure 4A and Figure 4B). In fact, when PC1 and PC2 scores were considered dependent variables and ‘species’ as independent variables, significant differences were observed between species (Wilcoxon non-parametric test: P < 0.01 for both PCs). Furthermore, the proportion of explained variance (R^2^) for PC1 and PC2 was 0.291 and 0.503, respectively. When the same analysis was repeated with ‘genomic compartments’ as an independent variable, significant differences were observed between compartments (Wilcoxon non-parametric test: P < 0.01 for both PCs), with R^2^ of 0.122 for PC1 and 0.335 for PC2. Consequently, when the first two PCs were considered, the proportion of the total variance explained by the ‘species’ and ‘compartments’ was 0.794 and 0.457, respectively. It was also evident that the genomic regions of *F. solani* were clearly distinguished from those of the other species, and that the genomic compartment B contained more heterogeneous regions than other compartments (Figure 4B).

**Figure 4.**
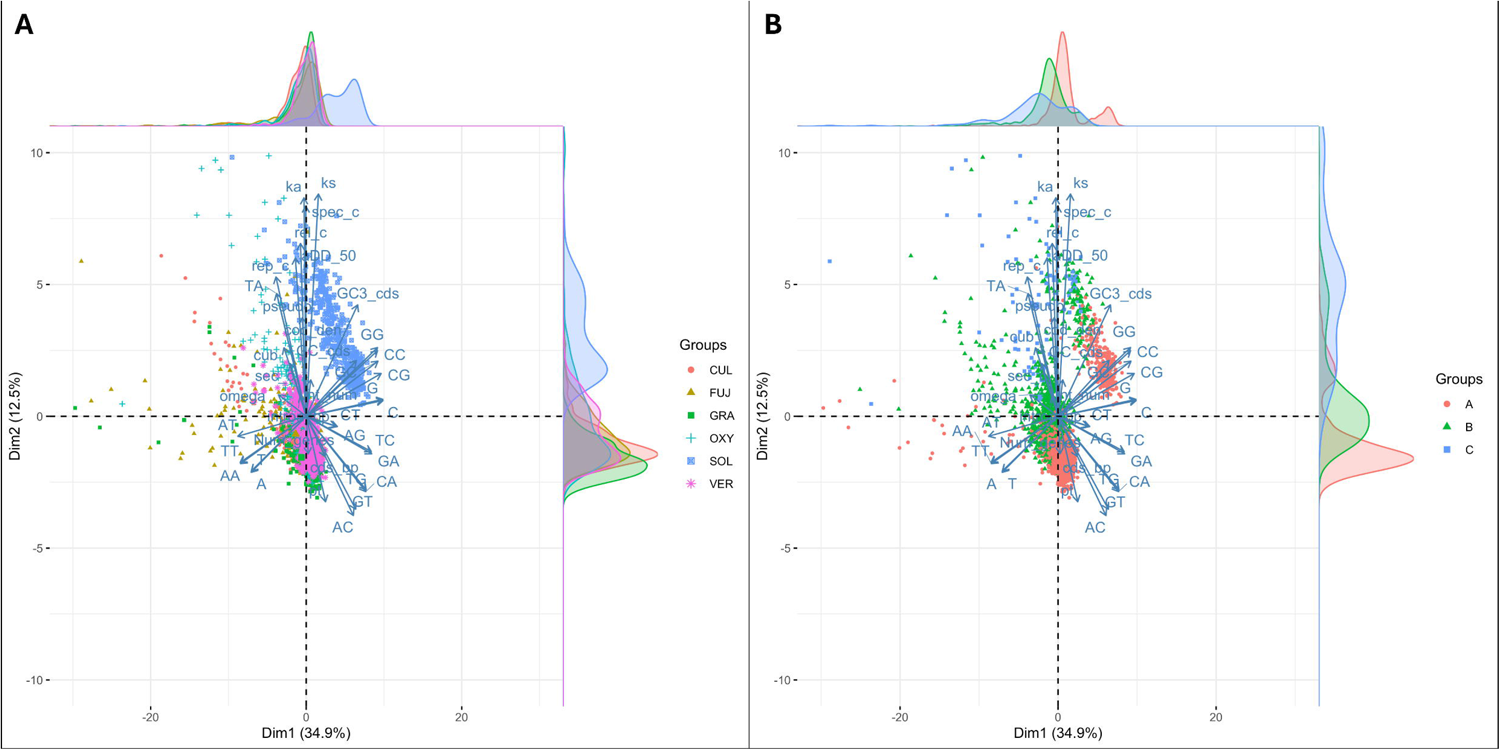
Principal component analysis (PCA) score plots depicting the relationship among all 100-kbp genomic regions of all *Fusarium* species based on all variables. Regions are marked according to the six investigated species (CUL for *Fusarium culmorum*; FUJ for *F. fujikoroi*; GRA for *F. graminearum*; OXY for *F. oxysporum*; SOL for *F. solani*; and VER for *F. verticilliodies*; panel A) or the compartment membership (compartment A, B and C; panel B).

Data were subjected to further analysis using canonical discriminant analysis, with all variables, to assign each genomic window to the species or compartment of origin (Figure S10 A-B). The frequency of misclassified genomic windows based on the first two canonical variables was 228 (9.56%) for species and 240 (10.07%) for compartments. Most misclassifications for species (165 out of 228) pertained to the pair *F. verticillioides* – *F. oxysporum*. Among the compartments, the highest proportion of correct assignments was observed for ‘A’, with 95.92% (48 misclassifications out of 1,423), followed by ‘B’ with 86.36% (111 misclassifications out of 814) and ‘C’ with 51.37% (71 misclassifications out of 146).

The variables describing the frequencies of mononucleotide and dinucleotide sequences showed the strongest correlation with PC1 and PC2, whereas genic variables loaded mainly on PC3 (data not shown).

Subsequently, two different PCAs were performed, one considering regional variables (Figure 5) and the other genic variables (Figure6).

**Figure 5.**
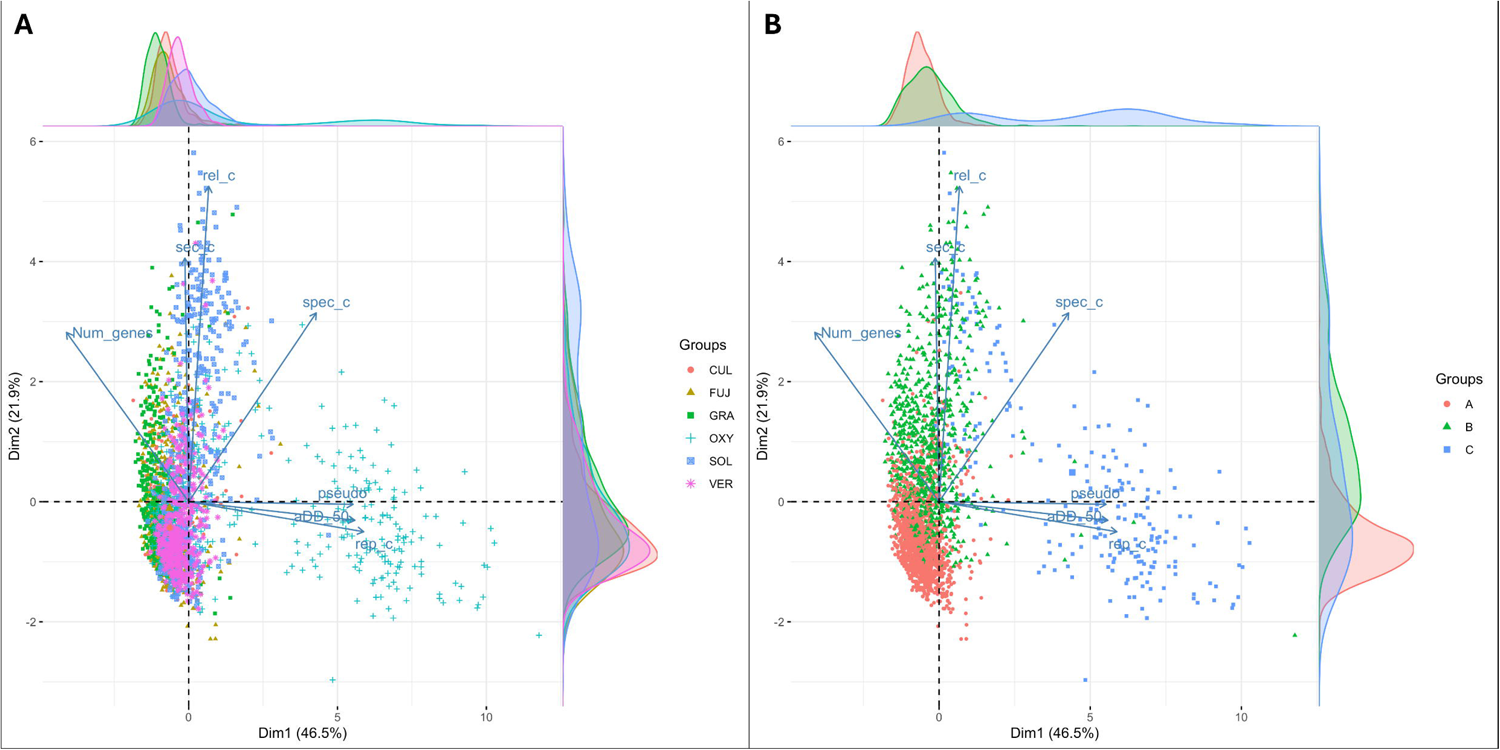
Principal component analysis (PCA) score plots depicting the relationship among all 100-kbp genomic regions of all *Fusarium* species based on regional structural variables. Regions are marked according to the six investigated species (CUL for *Fusarium culmorum*; FUJ for *F. fujikoroi*; GRA for *F. graminearum*; OXY for *F. oxysporum*; SOL for *F. solani*; and VER for *F. verticilliodies*; panel A) or the compartment membership (compartment A, B and C; panel B).

When only regional variables were considered, the regions of the different compartments were clearly separated (R^2^_PC1_ = 0.63 and R^2^_PC2_ = 0.33; Figure 5A) whereas the separation of the species was less clear (R^2^ _PC1_ = 0.33 and R^2^_PC2_ = 0-13). PC1 separated C from both compartments A and B, which differed mainly along PC2 (Figure 5B). The differentiation of genomic regions along PC1 was mainly driven by the density of pseudogenes, repetitive elements, and duplicated genes. Variables that explained the presence of species-specific, secreted, and relocated genes were the most important for the separation of the A and B compartments.

When only genic variables were considered (Figure 6), the genomic regions of compartment B were clearly separated from those of compartment A, but not from those of compartment C (R^2^_PC1_ = 0.54 and R^2^_PC2_ = 0.02; Figure 6B). The most significant variables in determining compartment differentiation were primary transcript length and non-synonymous substitution rates (Figure 6). It is important to note that the genomic regions of *F. solani* were not clearly distinguished from those of other species (Figure 6A). The genic variables were less effective in discriminating genomic regions according to the species along PC1 (R^2^_PC1_ = 0.27). The strong association between level of PC2 and species (R^2^_PC2_ = 0.65) was primarily attributable to the peculiarity of the *F.solani* genomic regions.

**Figure 6.**
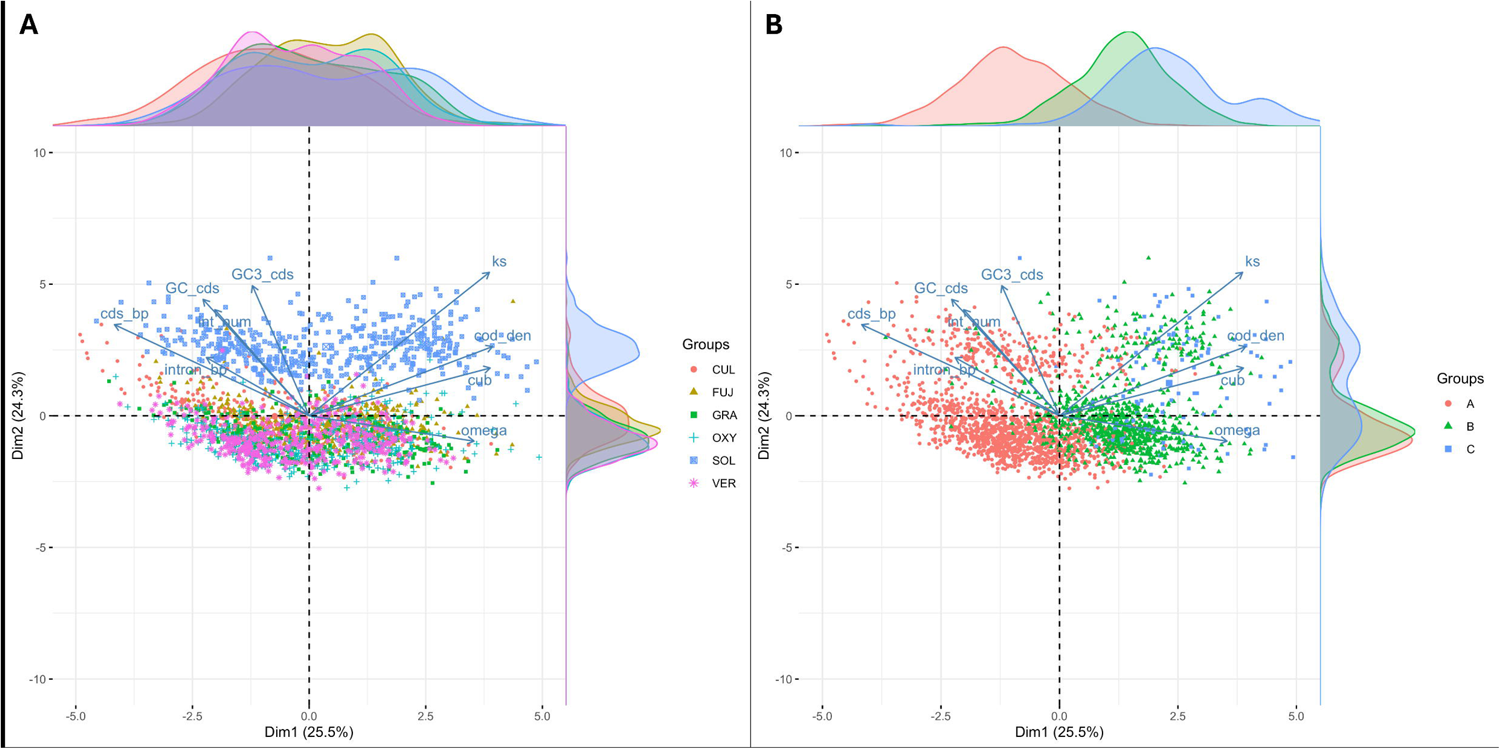
Principal component analysis (PCA) score plots depicting the relationship among all 100-kbp genomic regions of all *Fusarium* species based on regional variable only. Regions are marked according to the six investigated species (CUL for *Fusarium culmorum*; FUJ for *F. fujikoroi*; GRA for *F. graminearum*; OXY for *F. oxysporum*; SOL for *F. solani*; and VER for *F. verticilliodies*; panel A) or the compartment membership (compartment A, B and C; panel B).

The compositional regional variables (mononucleotide and dinucleotide regional content) were more effective in discriminating genomic regions based on species (R^2^_PC1_ = 0.24 and R^2^_PC2_ = 0.23) more efficiently than on compartment membership (R^2^_PC1_ = 0.11 and R^2^_PC 2_= 0.09)

The PCA performed on each species separately showed that, on average, the regions of compartments B and C were more closely related to each other than to those of compartment A (Figure 7). Another PCA was performed to examine the separation between the compartments B and C was conducted after excluding compartment A (Figure S7). The most pronounced differentiation was observed in the case of *F. oxysporum,* with DNA compositional variables exhibiting a strong correlated with PC1 and structural variables mainly correlated with PC2. Separation of the B and C compartments was less evident in *F. solani and F. fujikoroi*.

**Figure 7.**
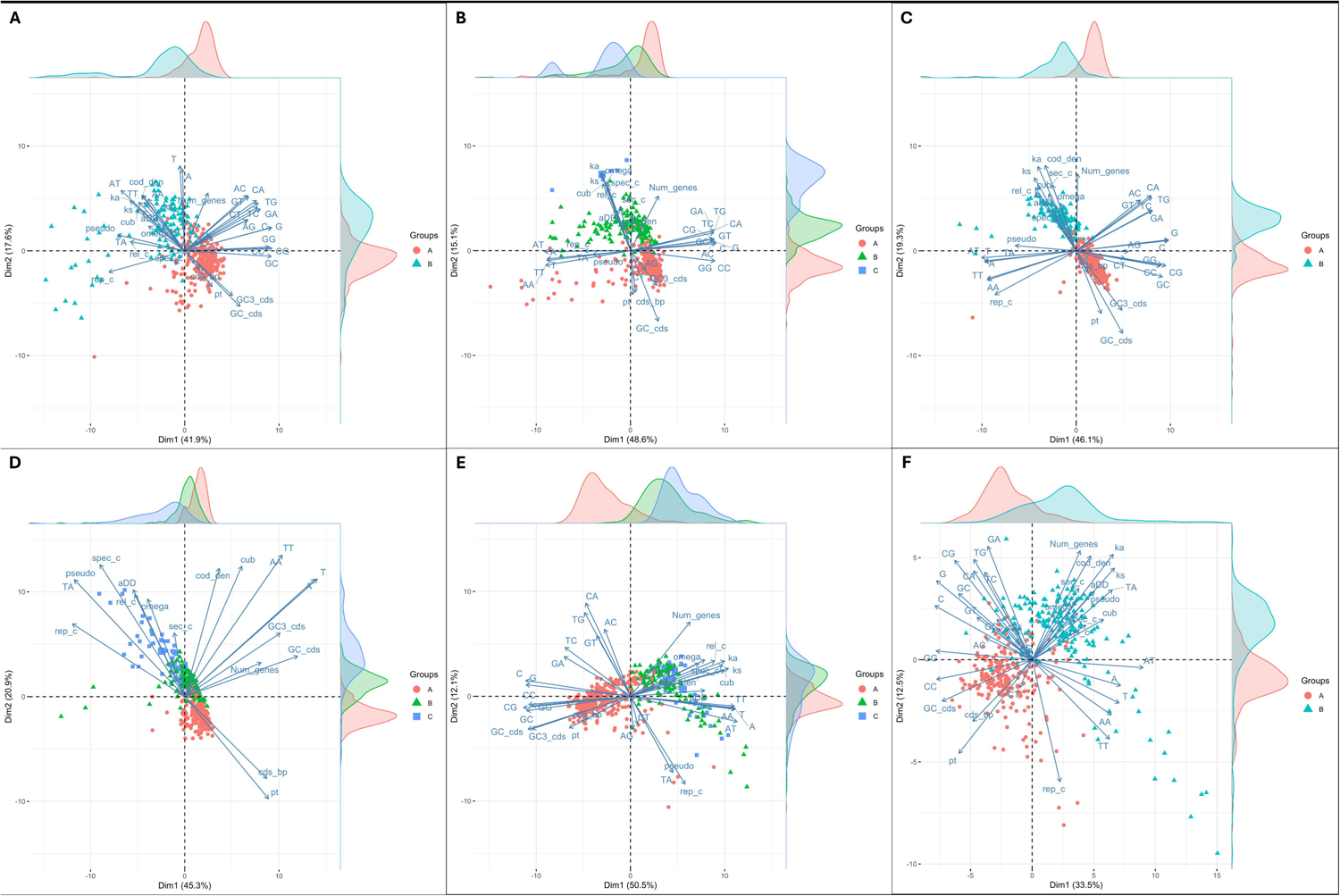
Principal component analysis (PCA) score plots depicting the relationship among the 100-kbp genomic regions for each of the six investigate *Fusarium* species based on all variables. Regions are marked according to the compartment membership (compartments A, B, and C): A) *Fusarium culmorum*, B) *F. Fujikoroi*, C) *F. graminearum*, D) *F. oxysporum*, E) *F. solani*, F) *F. verticillioides*.

## DISCUSSION

Genomic compartments were identified through the analysis of collinearity maps of six Fusarium species (Bertazzoni *et al*., 2018; Hane *et al*., 2011). Genomic sequences were divided into adjacent regions, which were then subjected to screening for structural and compositional variables. Significant associations between compartment membership and several of these variables were identified. Some of these associations, such as those with abundance of relocated and species-specific genes, were anticipated, given that disrupting collinearity has been shown to result in such patterns (Zhao et al., 2014). Other variables, such as those pertaining to DNA compositional and structural characteristics of genes (e.g., coding density or primary transcript length), are not obviously linked to collinearity and thus to compartment definition.

The observed diversity between genomic regions can be explained in terms of both species and compartment. The structural regional and genic variables accounted for the majority of the observed diversity due to compartment membership whereas the (regional) compositional variables exhibited greater variability among for species.

Consequently, although the majority of the events which generated the structural variability in genomes probably occurred after the radiation of the species, the resulting genomic compartments were similar, and the underlying mechanisms were probably also similar.

The DNA compositional variables (mono- and dinucleotide differences) were the primary contributors to species differentiation in all the analysed species. However, within each species the regional compositional variables were found to differentiate the genomic regions based on compartment membership. This indicates that the nucleotide compositional diversity of compartments is a common feature of Fusarium genomes. In general, regions with moderate or low ILC (i.e., less conserved across species) have a lower GC content than those with high ILC (i.e., highly conserved regions). However, the differences in dinucleotide frequency among compartments could not be explained by mononucleotide frequencies, suggesting the potential involvement of compositional bias in compartment evolution (Kariin and Burge, 1995; Porceddu and Camiolo, 2011).

In the yeast *Saccharomyces cerevisiae,* GC-rich isochores exhibit a chromatin structure that results in chromatin interactions with a lower frequency compared to A+T isochores (Dekker, 2007; Kiktev *et al*., 2018). A number of studies have indicated that genomic compartments, defined on the basis of collinearity, are associated with the structure of the chromatin in *Fusarium* spp. (Connolly *et al*., 2013; Bachleitner *et al*., 2021; Torres *et al*., 2020). Chromatin is a dynamic structure that influences the physical compaction and accessibility of DNA, thereby regulating several cellular functions, including chromosome segregation and gene transcription (Seidl *et al*., 2016; Strauss and Reyes-Dominguez, 2011; Elías-Villalobos *et al*., 2019; Buscaino, 2019). The clustering of genes, which are active only under specific conditions and in particular genomic compartments, allows chromatin-based regulation. In the absence of inducing conditions, the genomic region with densely compacted chromatin will be functionally repressed. Conversely, in the presence of inducing conditions, such as the presence of a compatible host or, for example, a medium with low nitrogen content, a change in chromatin compaction is favourable for gene expression (Connolly *et al*. 2013). Experimental evidence for such a mechanism has been provided in *F. graminearum* mutants lacking the activity of the kmt6 gene, which is responsible for the modification of H3K27me2 histone (Connolly et al. 2013). Modifications at H3K4me2/3 are associated with active chromatin, while H3K27me2 and H3K9me3 are associated with silent chromatin. A comparison of genome-wide histone modification and synteny maps revealed that H3K27me2 modifications overlapped with regions of low collinearity, while H3K3me2 modifications overlapped with regions of high collinearity. The disruption of the KMT6 gene results in the loss of H3K27me3 methylation and the derepression of the majority of genes found in regions of low collinearity. The majority of genes that are derepressed in the kmt6 knockout mutant are also derepressed in low nitrogen-containing medium, a condition that is known to induce the production of secondary metabolites and the expression of pathogenesis-related genes (Connolly *et al*., 2013). There was a minimal difference in expression between the wild-type strain and the kmt6 strain for primary metabolism genes indicating that the modification of H3K27me2 is the main responsible for the formation of facultative chromatin in *F. graminearum*. It has been demonstrated that facultative heterochromatin is associated with the modification of H3K27me2 as reported in *F. fujikoroi* and F. verticillioides (Connolly et al., 2013). Furthermore, the reduced expression of *F. fujikoroi* Kmt6 induced the expression of genes related to secondary metabolism and pathogenesis, in agreement with the observation carried out in *F. graminearum* (Connolly *et al*., 2013).

Regions with low or no collinearity were enriched in A- and T-containing oligonucleotides in a number of fungal species. Studies conducted in *F. circinatum* have indicated that repeat-induced point mutation (RIP) may result in the enrichment of A and T nucleotides, which could ultimately lead to the formation of facultative heterochromatin (Van Wyk *et al*., 2019; van Wyk *et al*., 2021). In *Neurospora crassa,* regions that have been identified as relics of RIP activity are methylated and targeted for incorporation into facultative heterochromatin, representing 4-6% of the genome (Lewis et al., 2009). Another noteworthy feature of these regions is the high density of pseudogenes (van Wyk et al.; Lewis et al., 2009). Our findings indicate that the association between the genomic compartment and the composition indices of the RIP is not consistent across species. It is notable that the C compartment of *F. oxysporum* exhibits a strong association, and that, in accordance with with expectations, its regions were also enriched for pseudogenes. However, in *F. oxysporum,* we found that the regions of compartment C were also enriched for duplicated and relocated genes, a finding apparently in contradiction with the RIP action, which is induced and targeted at duplicated sequences (Aramayo and Selker, 2013). Theoretical calculations indicate that RIP-induced mutations result in the rapid inactivation of genes. Consequently, the contribution of RIP to genic sub-functionalisation and neo-functionalisation is expected to be relatively modest (Galagan and Selker, 2004). However, if RIP activity is attenuated in one compartment, it could result in the production of mutated functional genes and, thereby significantly contributing to gene evolution. As elucidated by Galagan et al. (2004), if a gene is duplicated and one of the two copies is lost (deleted) by chance, the remaining copy will no longer be a substrate of RIP and will survive as a mutated copy of the original gene. This mechanism of transient duplication will result in a bias in the distribution of translocated genes (Galagan and Selker, 2004).

An intriguing observation of our work concerns the structural differences between genes residing in different compartments. In particular, our attention was directed to the exon-intron structure, which in our analysis was summarised by coding density (i.e. the ratio between the length of the coding exon and the length of the primary transcript). The findings of our study indicate that genes mapping in regions with low ILC exhibited higher coding densities than those in high ILC compartments in all the analysed species. A high coding density may be related to the effects of natural selection mediated by gene expression (Eisenberg and Levanon, 2003). As observed in numerous taxa, housekeeping genes have longer and more introns, while genes that are highly expressed have shorter and fewer introns to minimise the cost of transcription (Castillo-Davis *et al*., 2002; Camiolo *et al*., 2009). It is conceivable that genes that are expressed at high levels under inducing conditions also have an efficient structure that minimises the energetic costs of transcription. This could be of crucial importance in the context of responses to environmental stimuli, such as in the case of host-pathogen interaction. Nevertheless, the strength of such selection is relatively weak, and thus it should be evident only in species with large population sizes. Further studies are required to verify this hypothesis by examining the relationship between gene expression and structure with population size.

A genome-wide analysis of recombination in *F. graminearum* revealed that high recombination rates correspond to the B compartment (Laurent *et al*., 2017). This finding may help to explain some of the observations on genic structures. A high recombination rate reduces the linkage between sites and thus increases the effectiveness of selection, a phenomenon known as the Hill-Robertson effect (Comeron *et al*., 2007). It is conceivable that non-coding sequences, such as introns, may play a beneficial role in mitigating the intragenic Hill-Robertson effect. This is because they contribute to the spacing of selection targets, which are the coding regions. It was observed that genes located in compartment B (high recombination rate) of *F. graminearum* have a high coding density, and we observed that this is consistent across all the other species. It is therefore postulated that genes located in high recombination regions may be better able to cope with the Hill-Robertson effect, resulting in a more compact and energetically efficient structure. Conversely, genes mapped within compartment A experience a lower recombination rate and show a tendency to separate coding exons with long introns (Kreitman and Wayne, 1994) in order to mitigate the Hill-Robertson effect. Despite the differing absolute values observed, such the observed pattern was consistent across all the Fusarium species investigated.

Another potential explanation may be related to the different mutational biases acting on the gene compartments. Chen and Tyler (2022) have reported associations between the distribution of chromatin marks and the mechanism of DNA double-strand break repair have been reported. In particular, the H3K9me2/3 marks that are characteristic of heterochromatic regions have been associated with homologous recombination, whereas the facultative heterochromatin mark H3K27me has been associated with MMEJ (microhomology-mediated end joining), a double-strand break repair mechanism that is prone to introducing errors and short deletions. Consequently, the prevalence of MMEJ over other repair systems in compartments associated with facultative heterochromatin could explain the higher frequency of single nucleotide variants (SNVs) and the higher compactness of genic models (Ogata et al., 1996).

## References

Alexa, A., J. Rahnenführer, and T. Lengauer, 2006 Improved scoring of functional groups from gene expression data by decorrelating GO graph structure. Bioinformatics 22: 1600–1607.

Almagro Armenteros, J. J., K. D. Tsirigos, C. K. Sønderby, T. N. Petersen, O. Winther et al., 2019 SignalP 5.0 improves signal peptide predictions using deep neural networks. Nat Biotechnol 37: 420–423.

Altschul, S. F., W. Gish, W. Miller, E. W. Myers, and D. J. Lipman, 1990 Basic local alignment search tool. J Mol Biol 215: 403–410.

Aramayo, R., and E. U. Selker, 2013 Neurospora crassa, a model system for epigenetics research. Cold Spring Harb Perspect Biol 5.

Bertazzoni, S., A. H. Williams, D. A. Jones, R. A. Syme, K.-C. Tan et al., 2018 Accessories Make the Outfit: Accessory Chromosomes and Other Dispensable DNA Regions in Plant-Pathogenic Fungi. 31: 779–788.

Brown, N. A., J. Antoniw, and K. E. Hammond-Kosack, 2012 The Predicted Secretome of the Plant Pathogenic Fungus Fusarium graminearum: A Refined Comparative Analysis (S. Harris, Ed.). PLoS One 7: e33731.

Buscaino, A., 2019 Chromatin-Mediated Regulation of Genome Plasticity in Human Fungal Pathogens. Genes 2019, Vol. 10, Page 855 10: 855.

Camiolo, S., and A. Porceddu, 2013 gff2sequence, a new user friendly tool for the generation of genomic sequences. BioData Min 6.

Camiolo, S., D. Rau, and A. Porceddu, 2009 Mutational biases and selective forces shaping the structure of Arabidopsis genes. PLoS One 4.

Coleman, J. J., S. D. Rounsley, M. Rodriguez-Carres, A. Kuo, and C. C. Wasmann, 2009 The Genome of Nectria haematococca: Contribution of Supernumerary Chromosomes to Gene Expansion. PLoS Genetics | www 5: 1000618.

Connolly, L. R., K. M. Smith, and M. Freitag, 2013 The Fusarium graminearum Histone H3 K27 Methyltransferase KMT6 Regulates Development and Expression of Secondary Metabolite Gene Clusters (H. D. Madhani, Ed.). PLoS Genet 9: e1003916.

Depotter, J. R. L., X. Shi-Kunne, H. Missonnier, T. Liu, L. Faino et al., 2019 Dynamic virulence-related regions of the plant pathogenic fungus Verticillium dahliae display enhanced sequence conservation. Mol Ecol 28: 3482–3495.

Dong, S., S. Raffaele, and S. Kamoun, 2015 The two-speed genomes of filamentous pathogens: Waltz with plants. Curr Opin Genet Dev 35: 57–65.

Eisenberg, E., and E. Y. Levanon, 2003 Human housekeeping genes are compact. Trends Genet 19: 362–365.

Elías-Villalobos, A., R. R. Barrales, and J. I. Ibeas, 2019 Chromatin modification factors in plant pathogenic fungi: Insights from Ustilago maydis. Fungal Genetics and Biology 129: 52–64.

Emanuelsson, O., S. Brunak, G. von Heijne, and H. Nielsen, 2007 Locating proteins in the cell using TargetP, SignalP and related tools. Nat Protoc 2: 953–971.

Emms, D. M., and S. Kelly, 2019 OrthoFinder: Phylogenetic orthology inference for comparative genomics. Genome Biol 20: 1–14.

Faino, L., M. F. Seidl, X. Shi-Kunne, M. Pauper, G. C. M. van den Berg et al., 2016 Transposons passively and actively contribute to evolution of the two-speed genome of a fungal pathogen. Genome Res 26: 1091–1100.

Flynn, J. M., R. Hubley, C. Goubert, J. Rosen, A. G. Clark et al., 2020 RepeatModeler2 for automated genomic discovery of transposable element families. Proc Natl Acad Sci U S A 117: 9451–9457.

Frantzeskakis, L., B. Kracher, S. Kusch, M. Yoshikawa-Maekawa, S. Bauer et al., 2018 Signatures of host specialization and a recent transposable element burst in the dynamic one-speed genome of the fungal barley powdery mildew pathogen. BMC Genomics 19: 1–23.

Frantzeskakis, L., S. Kusch, and R. Panstruga, 2019 The need for speed: compartmentalized genome evolution in filamentous phytopathogens. Mol Plant Pathol 20: 3–7.

Galagan, J. E., and E. U. Selker, 2004 RIP: The evolutionary cost of genome defense. Trends in Genetics 20: 417–423.

Hart, M. L. I., B. L. Vu, Q. Bolden, K. T. Chen, C. L. Oakes et al., 2018a Genes Relocated Between Drosophila Chromosome Arms Evolve Under Relaxed Selective Constraints Relative to Non-Relocated Genes. J Mol Evol 86: 340–352.

Hart, M. L. I., B. L. Vu, Q. Bolden, K. T. Chen, C. L. Oakes et al., 2018b Genes Relocated Between Drosophila Chromosome Arms Evolve Under Relaxed Selective Constraints Relative to Non-Relocated Genes. J Mol Evol 86: 340–352.

Horton, P., K. J. Park, T. Obayashi, N. Fujita, H. Harada et al., 2007 WoLF PSORT: Protein localization predictor. Nucleic Acids Res 35: W585.

Kariin, S., and C. Burge, 1995 Dinucleotide relative abundance extremes: a genomic signature. Trends in Genetics 11: 283–290.

Karlin, S., 1998 Global dinucleotide signatures and analysis of genomic heterogeneity. Curr Opin Microbiol 1: 598–610.

Karlin, S., and I. Ladunga, 1994 Comparisons of eukaryotic genomic sequences. Proc Natl Acad Sci U S A 91: 12832–12836.

Karlin, S., and J. Mrázek, 1997 Compositional differences within and between eukaryotic genomes. Proc Natl Acad Sci U S A 94: 10227–10232.

Klee, E. W., and L. B. M. Ellis, 2005 Evaluating eukaryotic secreted protein prediction. BMC Bioinformatics 6.

Krogh, A., B. Larsson, G. Von Heijne, and E. L. L. Sonnhammer, 2001 Predicting transmembrane protein topology with a hidden Markov model: Application to complete genomes. J Mol Biol 305: 567–580.

Krzywinski, M., J. Schein, I. Birol, J. Connors, R. Gascoyne et al., 2009 Circos: An information aesthetic for comparative genomics. Genome Res 19: 1639–1645.

Lewis, Z. A., S. Honda, T. K. Khlafallah, J. K. Jeffress, M. Freitag et al., 2009 Relics of repeat-induced point mutation direct heterochromatin formation in Neurospora crassa. Genome Res 19: 427–437.

Löytynoja, A., and N. Goldman, 2010 WebPRANK: A phylogeny-aware multiple sequence aligner with interactive alignment browser. BMC Bioinformatics 11: 579.

Ma, L. J., H. C. Van Der Does, K. A. Borkovich, J. J. Coleman, M. J. Daboussi et al., 2010 Comparative genomics reveals mobile pathogenicity chromosomes in Fusarium. Nature 464: 367–373.

Ma, L.-J., D. M. Geiser, R. H. Proctor, A. P. Rooney, K. O’Donnell et al., 2013 Fusarium pathogenomics. Annu Rev Microbiol 67: 399–416.

Mandeel, Q., N. Ayub, and J. Gul, 2005 Survey of Fusarium species in an arid environment of Bahrain. VI.Biodiversity of the genus Fusarium in root-soil ecosystem of halophytic date palm (Phoenix dactylifera) community. 26: 365–404.

Mascagni, F., G. Usai, A. Cavallini, and A. Porceddu, 2021 Structural characterization and duplication modes of pseudogenes in plants. Sci Rep 11.

Novembre, J. A., 2002 Accounting for background nucleotide composition when measuring codon usage bias [2]. Mol Biol Evol 19: 1390–1394.

Ogata, H., W. Fujibuchi, and M. Kanehisa, 1996 The size differences among mammalian introns are due to the accumulation of small deletions. FEBS Lett. 390: 99–103.

Quinlan, A. R., and I. M. Hall, 2010 BEDTools: a flexible suite of utilities for comparing genomic features. Bioinformatics 26: 841–842.

R Core Team, 2013 R: A language and environment for statistical computing.

Seidl, M. F., D. E. Cook, and B. P. H. J. Thomma, 2016 Chromatin Biology Impacts Adaptive Evolution of Filamentous Plant Pathogens. PLoS Pathog 12.

Sievers, F., A. Wilm, D. Dineen, T. J. Gibson, K. Karplus et al., 2011 Fast, scalable generation of high-quality protein multiple sequence alignments using Clustal Omega. Mol Syst Biol 7: 539.

Slater, G. S. C., and E. Birney, 2005 Automated generation of heuristics for biological sequence comparison. BMC Bioinformatics 6: 31–31.

Smit, A., R. Hubley, and G. P, 2013 RepeatMasker Open-4.0.

Strauss, J., and Y. Reyes-Dominguez, 2011 Regulation of secondary metabolism by chromatin structure and epigenetic codes. Fungal Genet Biol 48: 62–69.

Torres, D. E., U. Oggenfuss, D. Croll, and M. F. Seidl, 2020 Genome evolution in fungal plant pathogens: looking beyond the two-speed genome model. Fungal Biol Rev 34: 136–143.

Visser, I., and M. Speekenbrink, 2010 depmixS4: An R package for hidden markov models. J Stat Softw 36: 1–21.

Wang, Y., H. Tang, J. D. Debarry, X. Tan, J. Li et al., 2012 MCScanX: A toolkit for detection and evolutionary analysis of gene synteny and collinearity. Nucleic Acids Res 40:.

Van Wyk, S., C. H. Harrison, B. D. Wingfield, L. De Vos, N. A. Van Der Merwe et al., 2019 The RIPper, a web-based tool for genome-wide quantification of Repeat-Induced Point (RIP) mutations. PeerJ 2019.

van Wyk, S., B. D. Wingfield, L. de Vos, N. A. van der Merwe, Q. C. Santana et al. Repeat-Induced Point Mutations Drive Divergence between Fusarium circinatum and Its Close Relatives.

Yang, Z., 2007 PAML 4: Phylogenetic analysis by maximum likelihood. Mol Biol Evol 24: 1586–1591.

Yang, H., H. Yu, and L. J. Ma, 2020 Accessory chromosomes in fusarium oxysporum. Phytopathology 110: 1488–1496.

Yu, G., F. Li, Y. Qin, X. Bo, Y. Wu et al., 2010 GOSemSim: an R package for measuring semantic similarity among GO terms and gene products. Bioinformatics 26: 976–978.

Zhang, Y., H. Yang, D. Turra, S. Zhou, D. H. Ayhan et al., 2020 The genome of opportunistic fungal pathogen Fusarium oxysporum carries a unique set of lineage-specific chromosomes. Commun Biol 3:

Zhao, C., C. Waalwijk, P. J. G. M. de Wit, D. Tang, and T. van der Lee, 2014 Relocation of genes generates non-conserved chromosomal segments in Fusarium graminearum that show distinct and co-regulated gene expression patterns. BMC Genomics 15: 191.

