## Supplementary Figures and Tables Captions for "Comparative analysis of the genomic architecture of six Fusarium species"

Supplementary material to “**Comparative analysis of genomic architecture of six Fusarium species”**

### Supplementary Table Captions

**Supplementary Table S1. Genomic compartments.** The genomic compartment of each genomic region is indicated as a letter (column 10). The coordinates of each region are indicated in column 1 (chromosome), 2 (region start) and 3 (region end). As a reference, the total number of genes mapping within each region is reported in column 4. Columns 5-9 indicate the number of genes that are collinear orthologous in the other species.
NOTE: CUL = *Fusarium culmorum*; FUJ = *F. fujikoroi*; GRA = *F. graminearum*; OXY = *F. oxysporum*; SOL = *F. solani*; VER = *F. verticilliodies*.

**Supplementary Table S2. Structural and compositional variables are associated with compartment membership.** Each genomic region was analysed for several structural variables. Differences between genomic compartments were analysed using the Wilcox test and a permutation approach. For each table are reported: the variable name (column 1), the species analysed (column 2), the genomic compartments (column 3-5), the *P-value* of the Wilcox test (column 6), the difference between each pair of compartments along with the values corresponding to the 1%, 5%, 95%, and 99% percentiles of the simulated variable distribution.

**Supplementary Table S3. The genomic compartments are differentiated for several structural and genomic features of genes mapping therein.** Genes were annotated for the presence of duplicate copies (column 5), the probability of the encoded protein to be secreted (column 6), the probability of being relocated (column 7) and the presence of homologous sequence in at least one of the cognate species (column 8).

**Supplementary Table S4. Distribution of regions scored as affected by Repeat induced point mutation among genomic compartments.** A 100-kb genomic region was defined as RIP-interested if it showed at least one subregion with a positive composite index value (Van Wyk *et al.*, 2019).

### Supplementary Figure captions

**Supplementary Figure S1. Trends for the Index of Local Collinearity (ILC) of *Fusarium culmorum* and the definition of genomic compartments.** For other details, please refer to the main text.

**Supplementary Figure S2. Trends for the Index of Local Collinearity (ILC) of *Fusarium fujikoroy* and the definition of genomic compartments.** For other details, please refer to the main text.

**Supplementary Figure S3. Trends for the Index of Local Collinearity (ILC) of *Fusarium solani* and definition of genomic compartments.** For other details, please refer to the main text.

**Supplementary Figure S4. Trends for the Index of Local Collinearity (ILC) of *Fusarium verticillioides* and definition of genomic compartments.** For other details, please refer to the main text.

**Supplementary Figure S5. Average gene densities in the genomic compartment of the *Fusarium* spp chromosomes.** The bar plots represent the average number of genes in the adjacent 100 kb regions of the genomic compartments of each chromosome. The compartments are distinguished by the colours of the boxes: grey = compartment A; brown = compartment B, and blue =compartment C.

**Supplementary Figure S6. Composition of orthologous groups.** Histograms represent the number of groups with at least one gene that belongs to the species indicated below as black dots. For example, 7408 groups were found to have at least one gene from the six species analysed (first bar on the histogram) whereas the number of groups with only *Fusarium solani* genes was 3200.

**Supplementary Figure S7. Mononucleotide relative content in in 100kb regions of the *Fusarium* spp genomic compartments.** The compartments are distinguished by the boxes' colours: blue = compartment C; green = compartment A, and red = compartment B.

**Supplementary Figure S8. Dinucleotide relative frequencies in 100kb regions of the genomic compartments of *Fusarium* spp.** The compartments are distinguished by the boxes' colours: blue = compartment C; green = compartment A, and red = compartment B.

**Supplementary Figure S9. Delta distances of Kariin (Kariin and Burge, 1995) between 100 kb genomic regions.** For each species, the distribution of delta distances among regions of different compartments (inter-compartment delta distances) is reported in the top graph. The distribution of delta distances between randomly sampled regions (reference delta distribution) is reported in the bottom graph. Wilcoxon tests applied to the delta distance between compartments versus the reference distributions were significant at $P < 0.01$ for all species except *F solani*.

**Supplementary Figure S10. Canonical discriminant analysis with all variables to assign each genomic window to the species (A) or the compartment of origin (B).** The abbreviations of species names are: CUL for *Fusarium culmorum*; FUJ for *F. fujikoroi*; GRA for *F. graminearum*; OXY for *F. oxysporum*; SOL for *F. solani*; and VER for *F. verticilliodies*.
Come è andatan
